# The L,D-transpeptidation pathway is inhibited by antibiotics of the β-lactam class in Clostridioides difficile

**DOI:** 10.1101/2024.06.13.598836

**Authors:** Ana M. Oliveira Paiva, Pascal Courtin, Glenn Charpentier, Olga Soutourina, Marie-Pierre Chapot-Chartier, Johann Peltier

## Abstract

The resistance of *Clostridioides difficile* to the β-lactam antibiotics cephalosporins, which target the peptidoglycan (PG) assembly, is a leading contributor to the development of *C. difficile* infections. *C. difficile* has an original PG structure with a predominance of 3→3 cross-links generated by L,D-transpeptidases (LDTs). *C. difficile* forms spores and we show that the spore cortex PG contains exclusively 3→3 cross-links. PG and spore cortex of *C. difficile* cells were largely unaffected by the deletion of the three predicted LDTs, revealing the implication of a new family of LDTS. The D,D-carboxypeptidases producing the essential LDT substrate were inactivated by cephalosporins, resulting in the inhibition of the L,D-transpeptidation pathway. In contrast, the participation of penicillin-binding proteins (PBPs) to PG cross-linking increased in the presence of the antibiotics. Our findings highlight that cephalosporin resistance is not primarily mediated by LDTs and illustrate the plasticity of the PG biosynthesis machinery in *C. difficile*.

## Introduction

*Clostridioides difficile* is the primary cause of antibiotic-associated nosocomial diarrhea in adults ^1^. *C. difficile* forms spores that are essential for the dissemination of *C. difficile* infections (CDI) ^2^. Antibiotic exposure resulting in gastrointestinal dysbiosis is commonly associated with the development of CDI ^1^. β-lactam antibiotics, particularly cephalosporins, are widely recognized as key factors in the development of CDI. However, the mechanism underlying the intrinsic resistance of *C. difficile* to these antibiotics remains poorly understood ^3,4^. β-lactams target the cell wall peptidoglycan (PG) assembly. PG is a polymeric macromolecule consisting of linear glycan chains cross-linked by short peptides ^5^. The D,D-transpeptidase (DDT) activity of penicillin-binding proteins (PBPs) connects the fourth amino acid of one peptide side chain to the third amino acid of another, forming 4→3 cross-links. However, 3→3 cross-links generated by L,D-transpeptidases (LDTs) presenting a YkuD-like domain have also been reported ^6^. In addition to their cross-linking activity, both DDTs and LDTs catalyze exchange reactions where the terminal D-Ala in pentapeptides and tetrapeptides, respectively, are replaced by a non-canonical D-amino acid (NCDAA) ^7,8^. DDTs, which are the primary targets of β-lactam antibiotics, use native PG precursors containing a pentapeptide stem as acyl donors for cross-linking. In contrast, LDT activity requires a tetrapeptide stem as the acyl donor substrate ^6^. This substrate is typically produced by D,D-carboxypeptidases that cleave the fifth residue from the pentapeptide chain ^9–11^. LDTs have the potential to confer resistance to β-lactam antibiotics, as they are solely inhibited by β-lactam antibiotics of the carbapenem class^12^.

*C. difficile* is the only known Bacillota with predominant 3→3 PG cross-links ^6,13,14^. Recent studies revealed that a mutant strain of *C. difficile* R20291, which lacks all three LDT paralogues, is not impaired in the production of 3→3 cross-links ^15,16^. Furthermore, two new enzymes with a VanW catalytic domain have been identified as the missing LDTs ^16^. However, the role of 3→3 cross-linking in mediating cephalosporin resistance in *C. difficile* remains unclear. While the structure of the *C. difficile* spore cortex, a modified layer of PG, has also been characterized, the type of cross-links present has not been defined ^17^.

In this study, we showed that the spore cortex PG of *C. difficile* only contains 3→3 cross-links. As for the PG of vegetative cells, 3→3 cross-links were still present in the spore cortex of a *C. difficile* strain lacking the three YkuD domain-containing LDTs, suggesting the implication of the VanW domain proteins. By treating *C. difficile* cells with subinhibitory concentrations of β-lactams, including cephalosporins and carbapenems, we discovered that at least one D,D-carboxypeptidase, which is necessary for the production of LDT substrates, is the primary target of these antibiotics and is differentially sensitive to them. In contrast, PBPs generating the 4→3 cross-links are not inhibited by these concentrations of antibiotics. Thus, cephalosporin resistance in *C. difficile* is not dependent on PG cross-linking by LDTs and instead relies on the poor inhibition of PBPs by this antibiotic family. These findings suggest that the growth arrest induced by β-lactams is mediated by the inhibition of the L,D-transpeptidation pathway in *C. difficile*.

## Results

### A strain lacking the three YkuD proteins still predominantly produces 3→3 cross-links

To evaluate the role of the three proteins harboring the catalytic domain YkuD on the PG structure of *C. difficile* 630Δ*erm*, we generated a triple mutant of the corresponding genes *ldt_Cd1_* (*CD2963*), *ldt_Cd2_* (*CD2713*) and *ldt_Cd3_* (*CD3007*)^14^ (Figures S1A and S1B). This mutant, hereby referred as ΔΔΔ*ldt*, was confirmed via whole-genome sequencing and no additional mutation could be detected (see STAR methods). The growth rate of the ΔΔΔ*ldt* strain was similar to that of the parental strain 630Δ*erm* (Figure S1C).

Wild-type and ΔΔΔ*ldt* cells were collected in exponential growth and the purified PG was digested with mutanolysin to generate muropeptides. Muropeptides were analyzed by ultra-high performance liquid chromatography coupled to tandem mass spectrometry (UHPLC-MS/MS) (Figure 1A) and their structure was deduced from their *m/z* values and fragmentation patterns obtained by MS/MS. The structure of all identified muropeptides is summarized in Table S1. The muropeptide composition of the ΔΔΔ*ldt* strain was largely similar to that of the parental strain (Figure 1A and Table S1). The main difference was a strong reduction of the abundance of muropeptides containing a tetrapeptide stem ending in Gly (peaks 4 and 11) in the mutant compared to the wild-type strain (Figures 1A and 1C and Table S1), revealing a role of at least one of the three enzymes in the exchange reaction of the terminal D-Ala in tetrapeptide stems with Gly. Additionally, a small but significant decrease of the dimers cross-linked by L,D-transpeptidation, which results in a reduction of the cross-linking index was observed in the ΔΔΔ*ldt* strain compared to the parental strain (Figures 1C and 1D). Nonetheless, 3→3 cross-links remained predominant in the triple mutant strain, indicating the presence of LDTs with a different catalytic domain compensating for the loss of the canonical LDTs.

**Figure 1.**
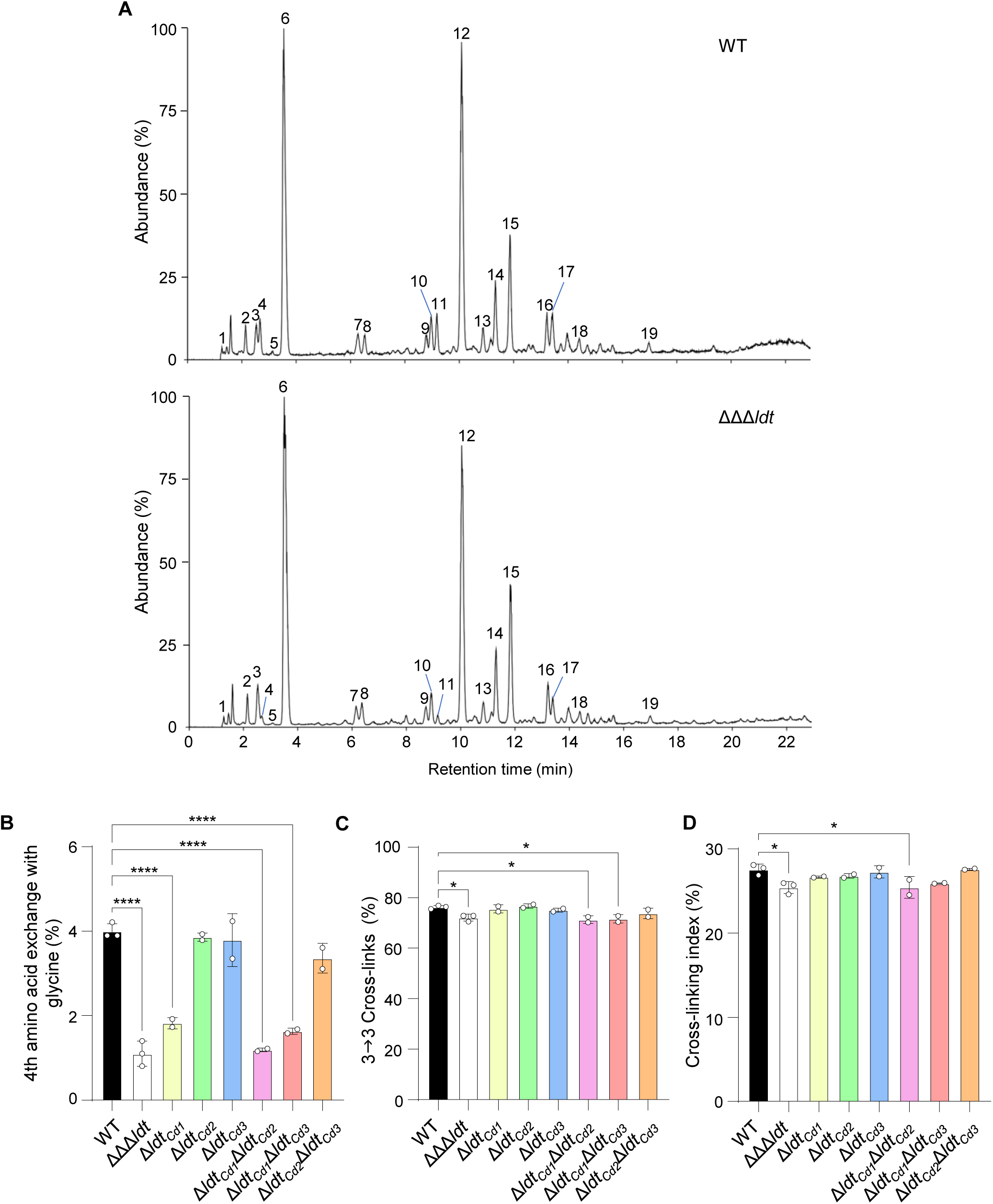
Impact of the *ldt* single, double and triple deletions on the PG structure of *C. difficile* vegetative cells. (A) LC-MS chromatograms of muropeptides from vegetative cells of *C. difficile* 630Δ*erm* wild-type (WT) and ΔΔΔ*ldt* strains. Major peaks are labelled with numbers referring to Table S1. See also Table S1 for the structure of all identified muropeptides. Data are representative of three independent experiments. (B) Abundance of muropeptides containing a modified tetrapeptide stem with the LDT-mediated exchange of the terminal D-Ala by a Gly in the PG from *C. difficile* WT and *ldt* mutant strains. (C) Abundance of muropeptide dimers with a 3→3 crosslink relative to the total cross-links in dimers in the PG from *C. difficile* WT and *ldt* mutant strains. (D) Cross-linking index of PG from *C. difficile* WT and *ldt* mutant strains. The cross-linking index was calculated as described by Glauner ^34^. All graphs represent mean ± SD and include individual data points; *n* = 3 independent experiments for WT and ΔΔΔ*ldt* and *n* = 2 independent experiments for the other mutant strains. \**P* ≤ 0.05 and \*\*\*\**P* ≤ 0.0001 by a one-way ANOVA followed by a Dunnett’s multiple comparisons test.

To confirm the impact of the mutations on the exchange reaction, *C. difficile* cells were labeled with the amino-acid probe HCC-amino D-alanine (HADA) ^18,19^ during exponential growth and visualized using fluorescence microscopy (Figure S2A). The cell length and width between the triple mutant and the parental strain were identical (Figures S2B and S2C). In pre-divisional cells, the intensity of the HADA staining significantly increased at mid-cell, as expected for the septum formation (Figures S2A, S2D and S2E). In dividing cells, a gradient of fluorescence was observed with a concentrated signal at one of the poles, which presumably corresponds to the newly formed pole, and a fading signal at the other pole (Figures S2A and S2E). The triple mutant showed a similar fluorescent labelling profile to the wild type around the cells, but an overall significant decrease in fluorescence intensity (Figures S2A and S2D). This result suggests that HADA incorporation through the exchange reaction into the PG of *C. difficile* is partially dependent on the activity of at least one of the LDTs.

### Ldt_Cd1_ contributes to the exchange reaction and the generation of 3→3 cross-links

To elucidate the contribution of the three LDTs to exchange reactions and 3→3 crosslinking, the muropeptide profile of the different single and double mutants was also analyzed (Figure S3 and Table S1). The abundance of tetrapeptides ending in Gly and of 3→3 cross-links in the strains deleted of *ldt_Cd2_* and/or *ldt_Cd3_* were similar to those of wild type (Figures 1B and 1C). However, deletion of either *ldt_Cd1_*, *ldt_Cd1_* and *ldt_Cd2_* or *ldt_Cd1_* and *ldt_Cd3_* recapitulated the decrease of tetrapeptides ending in Gly observed in the ΔΔΔ*ldt* strain. The proportion of 3→3 cross-links was also reduced in the Δ*ldt_Cd1_*Δ*ldt_Cd2_* and Δ*ldt_Cd1_*Δ*ldt_Cd3_* double mutants with an impact on the cross-linking index (Figures 1C and 1D). Thus, these data indicate that the PG modifications observed in the triple mutant strain are mostly mediated by the *ldt_cd1_* deletion.

### The spore cortex of *C. difficile* is cross-linked by L,D-transpeptidation

To determine the mode of cross-linking in the spore cortex, spores of *C. difficile* 630 Δ*erm* wild type were purified and their PG cortex was isolated. Muropeptides were generated using mutanolysin, analyzed by UHPLC-MS/MS (Figure 2A), and their structure was deduced from their *m/z* values and fragmentation patterns obtained by MS/MS. All the identified peaks are summarized in Table S2. In agreement with a previous report ^17^, 39.5% ± 3.7% of the NAG residues were found to be *N*-deacetylated in the spore cortex of *C. difficile* wild type (Figure 2B). In addition, 31.1 ± 2.9% of the NAM residues were converted to the spore-specific MAL in *C. difficile* spore cortex (Figure 2C). This modification prevents the cleavage between the NAM and NAG residues by mutanolysin, leading to the presence of oligosaccharides among the generated muropeptides. Most of the muropeptides still containing an unmodified NAM residue lacked any stem peptide (34.7 ± 3.1%) or harbored a tetrapeptide stem (30.6 ± 4.6%) (Figure 2C).

**Figure 2.**
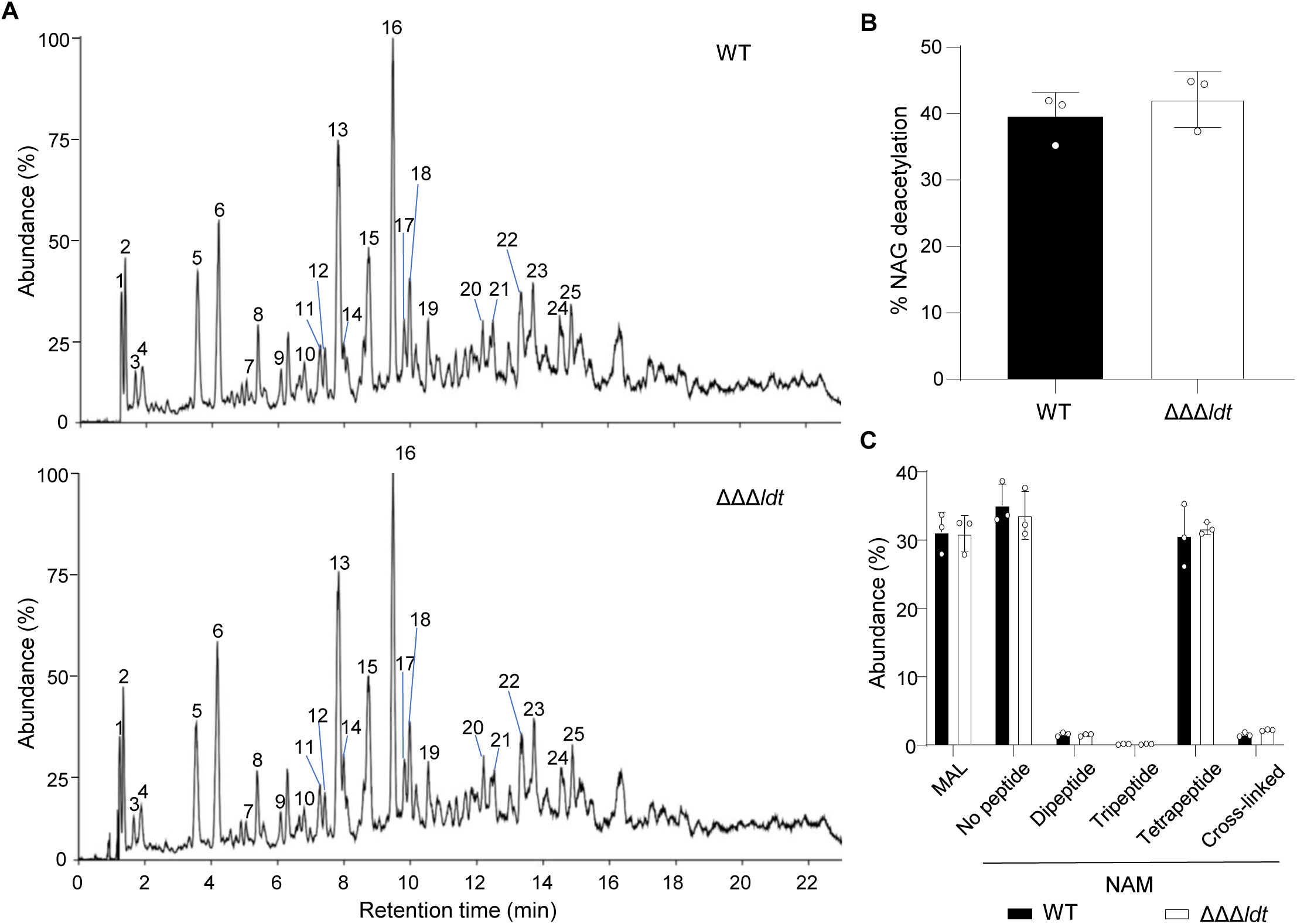
Impact of the *ldt* triple deletion on the PG structure of *C. difficile* spore cortex. (A) LC-MS chromatograms of muropeptides from the spore cortex of *C. difficile* 630Δ*erm* wild-type (WT) and ΔΔΔ*ldt* strains. Major peaks are labelled with numbers referring to Table S2. See also Table S2 for the structure of all identified muropeptides. Data are representative of three independent experiments. (B) Abundance of NAG deacetylation in the spore cortex from *C. difficile* WT and ΔΔΔ*ldt* strains. (C) Abundance of MAL and NAM in the spore cortex from *C. difficile* WT and ΔΔΔ*ldt* strains. Percentages of unsubstituted NAM (no peptide) and of NAM substituted with a dipeptide, a tripeptide or a tetrapeptide, or involved in cross-linking are represented. All graphs represent mean ± SD and include individual data points; *n* = 3 independent experiments.

Among the identified muropeptides, only two corresponded to dimers and no trimer was detected (Table S2). Thus, the spore cortex of *C. difficile* is very weakly cross-linked with a cross-linking index of 1.5 ± 0.3% (Figure 2C). The two dimers had the same structure with a tripeptide and a tetrapeptide stem and both contained a 3→3 cross-link as revealed by the loss of one C-terminal alanine (mass loss of 89.048 Da) from the acceptor tetrapeptide stem in their MS-MS fragmentation spectrum (Figures 3 and S4). Thus, the spore cortex of *C. difficile* contains only 3→3 cross-links.

**Figure 3.**
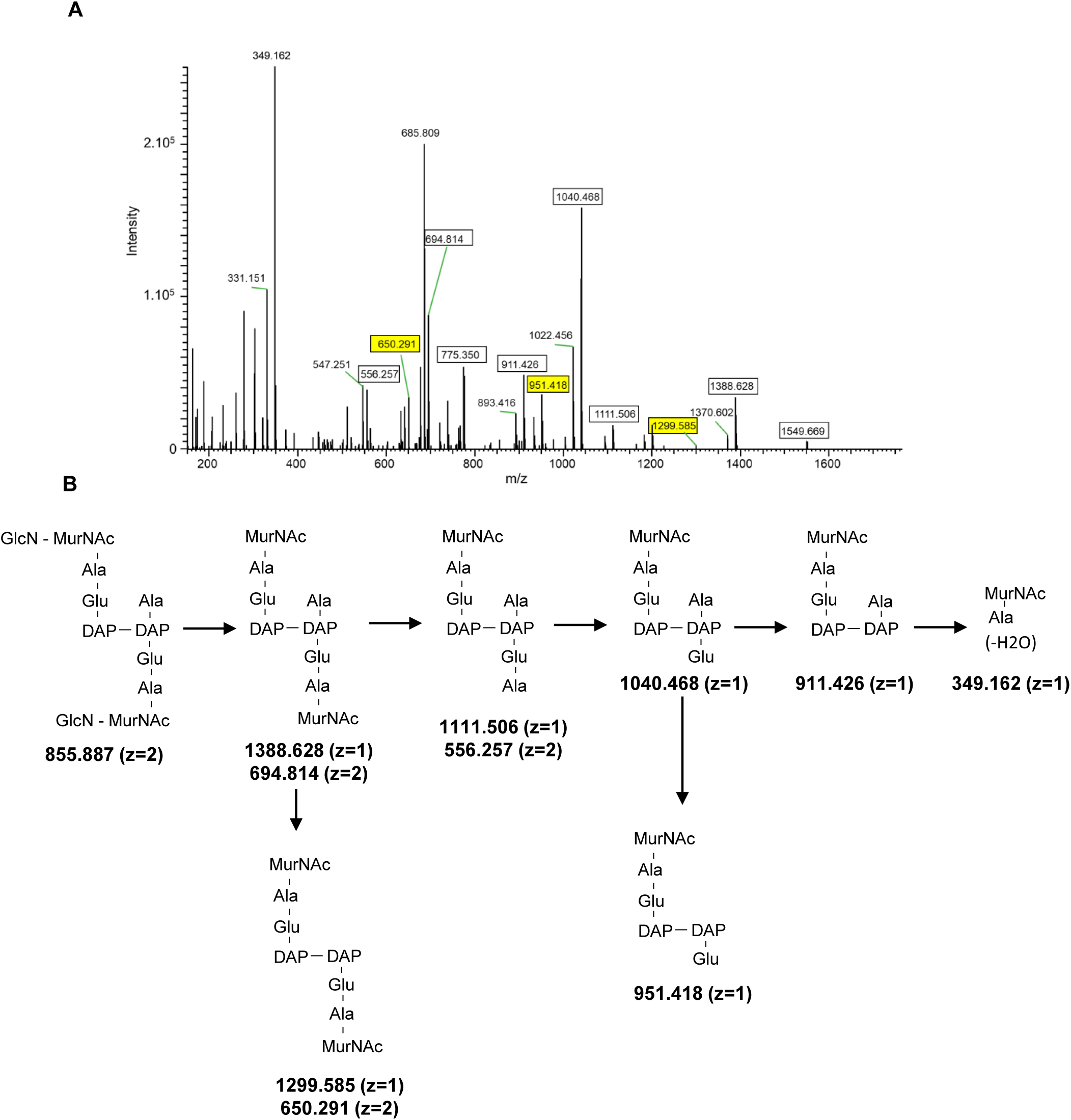
MS/MS analysis of the main muropeptide dimer from the spore cortex of *C. difficile* wild type. (A) MS/MS analysis of the molecular ion [M+2H]^2+^ (*m/z* = 855.887 with *z*=2) detected in peak 14 with a retention time of 7.99 min in Figure 4A and Table S2. Fragments (squared) were detected as [M+H]^+^ (z=1) or [M+2H]^2+^ (z=2) adducts. Fragments resulting from loss of a water molecule are not squared. The fragments allowing assignment of the 3-3 cross-links are highlighted in yellow. (B) Structures inferred from the MS/MS analysis are presented. The loss of one alanine from the C-terminal end of different ions (mass loss of 89.048) establishes the presence of a 3→3 crosslink.

Next, we analyzed the spore PG structure of the ΔΔΔ*ldt* strain to assess the implication of the LDTs in the formation of the 3→3 cross-links. However, the muropeptide profile of the mutant was identical to that of the wild-type strain and no decrease of the 3→3 cross-links was observed (Figure 2 and Table S2). These data indicate that either the canonical LDTS do not contribute to the cross-linking of the spore cortex or their loss is compensated by the activity of at least one of the two novel LDTs with a VanW domain ^16^.

### The YkuD domain proteins do not contribute to β-lactam resistance in *C. difficile*

We investigated the effect of the triple *ldt* deletion on β-lactam resistance in *C. difficile*. Loss of the three L,D-transpeptidases did not alter the minimum inhibitory concentration (MIC) of different members of the β-lactam antibiotics class (Table S3). In addition, growth of the ΔΔΔ*ldt* and the parental strains in the presence of subinhibitory concentrations of different β-lactams (1/2 MIC) were similar (Figure S5). We reasoned that the gene expression of the VanW domain LDTs or other enzymes of the PG polymerisation machinery might be induced in the presence of β-lactams to compensate for the loss of the three YkuD proteins. To test this hypothesis, 630Δ*erm* and ΔΔΔ*ldt* were grown in the presence of subinhibitory concentrations of the cephalosporin antibiotic ceftriaxone (1/2 MIC) and the samples were subjected to RNA-sequencing. Whereas genes involved in stress response, such as type I toxin-antitoxin systems were induced in ΔΔΔ*ldt*, no gene related to PG metabolism, including the VanW proteins (*CD1436* and *CD2149* in 630Δ*erm*) or encoding PG-associated proteins could be identified as differently expressed (Table S4). These data indicate that the YkuD domain proteins are dispensable for β-lactam resistance in *C. difficile* 630Δ*erm*.

### The L,D-transpeptidation pathway is inhibited by subinhibitory concentrations of cephalosporin and carbapenem antibiotics in *C. difficile*

To examine the impact of β-lactams on PG cross-linking, the PG structure of vegetative cells of the ΔΔΔ*ldt* strain grown in the presence of 1/2 MIC of the cephalosporin cefoxitin was analysed. *C. difficile* exposure to this antibiotic led to drastic PG structure modifications, in comparison to the non-treated strain, with the appearance of new abundant muropeptides with a pentapeptide stem to the detriment of tetrapeptide stems (Figures 1A, 4A and 4B and Table S1). Muropeptides with a pentapeptide stem accounted for 31.6 ± 2.4% of the total muropeptides in the cefoxitin-treated strain when they represented only 0.5 ± 0.04% in the untreated strain. Conversely, the abundance of tetrapeptides decreased from 70.5 ± 0.9% to 36.6 ± 0.7% in the presence of the antibiotic (Figure 4B). These changes were accompanied by a 50% decrease of the abundance of muropeptide dimers containing 3→3 cross-links (Figure 4C and Table S1). Remarkably, the cross-linking index remained similar in the absence and in the presence of antibiotics as the 3→3 cross-link reduction was compensated by an equivalent increase of the 4→3 cross-linked dimers (Figure 4B). Treatment with ceftriaxone, another antibiotic of the cephalosporin class, resulted in similar PG modifications, although at a lesser extent (Figure S6). Thus, these data strongly suggest that cephalosporins inactivate at high concentrations one or several D,D-carboxypeptidase(s), whose activity is required to produce the substrate of the LDT. They also reveal that at least one PBP catalysing the formation of the 4→3 cross-linking reaction is not inactivated by these antibiotics.

**Figure 4.**
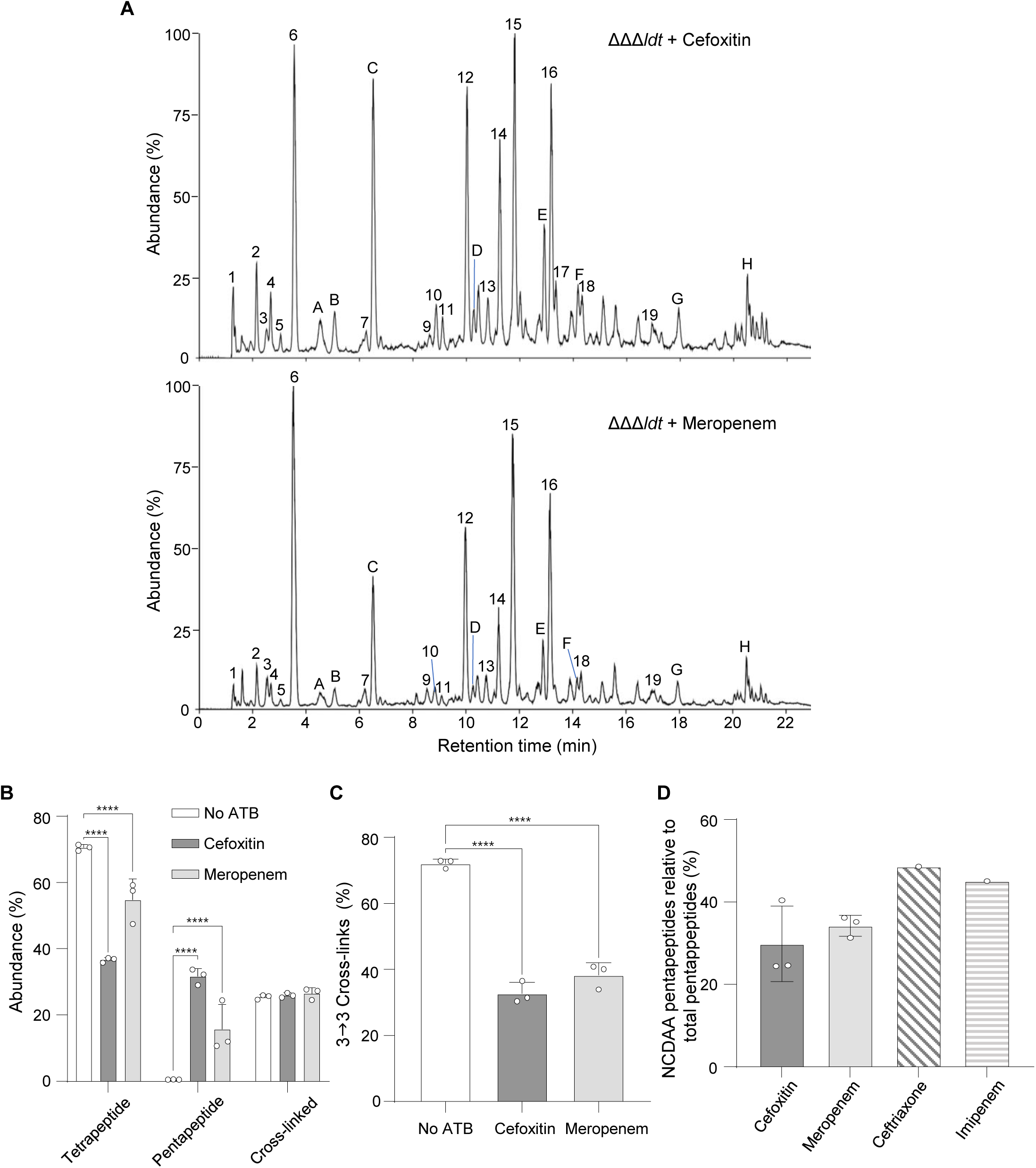
Impact of different β-lactam antibiotics on the PG structure of vegetative cells of *C. difficile* ΔΔΔ*ldt*. (A) LC-MS chromatogram of muropeptides from vegetative cells of *C. difficile* ΔΔΔ*ldt* strain grown in the presence of subinhibitory concentrations of cefoxitin or meropenem. Peak labels refer to Table S1. New major peaks observed in the presence of the antibiotics are labelled with letters. See also Table S1 for the structure of all identified muropeptides. Data are representative of three independent experiments. (B) Abundance of muropeptides with a tetrapeptide or a pentapeptide stem and cross-linking index in the PG of *C. difficile* ΔΔΔ*ldt* strain grown in the absence of antibiotics (no ATB) or in the presence of cefoxitin or meropenem. Means and SD are shown; *n* = 3 independent experiments. \*\*\*\**P* ≤ 0.001 by a two-way ANOVA followed by a Dunnett’s multiple comparisons test comparing values with the average value of the no ATB condition. (C) Abundance of muropeptide dimers with a 3→3 crosslink relative to the total cross-links in dimers in the PG of *C. difficile* ΔΔΔ*ldt* strain grown in the absence of antibiotics (no ATB) or in the presence of cefoxitin or meropenem. Means and SD are shown; *n* = 3 independent experiments. \*\*\*\**P* ≤ 0.001 by a one-way ANOVA followed by a Dunnett’s multiple comparisons test comparing values with the average value of the no ATB condition. (D) Abundance of muropeptides with a pentapeptide stem ending with a non-canonical D-amino-acid (NCDAA: Gly, Phe, Leu or Val) relative to the total pentapeptide muropeptides in the ΔΔΔ*ldt* strain. All graphs represent mean ± SD and include individual data points; *n* = 3 independent experiments. \*\*\*\**P* ≤ 0.001 by a two-way ANOVA followed by a Dunnett’s multiple comparisons test (B and C).

*C. difficile* 630Δ*erm* is resistant to most cephalosporins but susceptible to carbapenems (Table S3) ^20^. To assess whether carbapenems also impacted the activity of D,D-carboxypeptidases in *C. difficile*, PG structure of ΔΔΔ*ldt* cells treated with 1/2 MIC of meropenem or imipenem was analysed (Figures 1A and S6A and Table S1). As with cephalosporins, both carbapenem antibiotics induced the appearance of several major muropeptides peaks with pentapeptide stems and modified the balance between 4→3 and 3→3 cross-links to the benefits of 4→3 cross-links (Figures 4B, 4C, S6B and S6C). Similar results were obtained when analysing the PG structure of the wild-type strain grown in the presence of cefoxitin or meropenem ruling out an impact of the *ldt* deletions on the D,D-carboxypeptidase inactivation (Figures S7A, S7B and S7C).

Another remarkable effect of the antibiotic treatments on the PG structure was the presence of a high proportion of muropeptides with a pentapeptide stem ending with a non-canonical amino-acid (NCDAA: Gly, Phe, Leu or Val) (Table S1, Figures 4D and S7D). The incorporation of NCDAA into the PG was shown to regulate the cell wall structure in response to environmental stresses in *Vibrio cholerae* and the same mechanism was suggested in *Bacilus subtilis* ^8,21^.

Altogether, these data strongly suggest that at least one D,D-carboxypeptidase plays a pivotal role in the control of the mode of transpeptidation in *C. difficile*. This D,D-carboxypeptidase is the primary target of antibiotics of the β-lactam family and is differentially susceptible to these antibiotics with an efficient inhibition by low concentrations of carbapenems but only by high concentrations of cephalosporins. The associated reduction of 3→3 cross-linking, resulting from the limited LDT substrate availability, highlights that cephalosporin resistance is not primarily mediated by LDTs in *C. difficile* but rather relies on PBPs resistant to this class of antibiotics.

## Discussion

In agreement with recent studies ^15,16^, we showed here that a *C. difficile* 630Δ*erm* strain lacking the three predicted LDTs presented only a slight reduction of 3→3 cross-links and a decrease in the exchange reaction. The two novel LDTs with a VanW domain most likely compensate for the loss of the other enzymes as recently shown in the strain R20291 ^16^. PG analysis of the different *ldt* single and double mutant strains further identified Ldt_Cd1_ as the enzyme mediating most of the observed changes. This result is contrasting with *in-vitro* assays of the three recombinant LDTs showing that Ldt_Cd1_ is poorly active and has a weak activity in catalysis of the exchange reaction in comparison to Ldt_Cd2_ and Ldt_Cd3 15,22_.

We showed the abundant presence of muramic-δ-lactams and *N*-acetylmuramic acids with no peptide stem, which results in a low amount of cross-links, in the spore cortex of *C. difficile*. Furthermore, we discovered that 3→3 cross-links are largely predominant, if not exclusive, in the spore cortex. In *B. subtilis*, the enzymes required for the cross-linking of the spore cortex are produced in the mother cell and their expression is under control of SigE ^23^, a sporulation-specific sigma factor. Previous transcriptomic analyses identified *ldt_Cd3_* gene as a member of the SigE regulon in *C. difficile* ^24–26^. In addition, Ldt_Cd3_ was shown to display a strong LDT activity *in vitro* ^15,22^. Yet, deletion of the three LDT-encoding genes had no impact on the abundance of the cross-links in the cortex PG, implying the involvement of the non-canonical LDTs. Future work will be required to identify which LDTs contribute to the formation of the spore cortex peptidoglycan.

Among the inhibitors of PG polymerisation, β-lactam antibiotics stand out as the most clinically significant. The β-lactam ring found in all β-lactam antibiotics interacts with the essential nucleophilic serine of PBPs, which can display DDT, endopeptidase or D,D-carboxypeptidase activity ^27^. This interaction results in the creation of a stable acyl-enzyme adduct that interferes with catalysis. In contrast, LDTs are slowly acylated by β-lactams of the penicillin and cephalosporin classes and the resulting acyl-enzymes are rapidly hydrolysed ^28^. Thus, LDTs are insensitive to these antibiotics but are still efficiently inactivated by carbapenems ^29,30^. In agreement, activation of the L,D-transpeptidation pathway in *Enterococcus faecium* and *Escherichia coli* conferred broad-spectrum resistance to β-lactams, with the exception of carbapenems ^29,31^. Conversely, we showed that this pathway is inhibited by both cephalosporins and carbapenems in *C. difficile*, revealing that it does not mediate resistance to these drugs. An inhibition of the DDT activity of PBPs could explain the accumulation of muropeptides with a pentapeptide stem observed in the presence of the antibiotics. However, this hypothesis is not supported by our data as 4→3 PG cross-links were more abundant in antibiotic-treated cells. In line with this result, biochemical analyses of the purified class A PBP PBP1 and the class B PBP PBP2 of *C. difficile*, which are respectively essential for cross-links synthesis during vegetative cell division and elongation, revealed that both enzymes are insensitive to most cephalosporins and carbapenems ^32,33^ The only other explanation to the observed structural changes is the inhibition of at least one D,D-carboxypeptidase. Activation of the L,D-transpeptidation pathway leading to β-lactam resistance in *E. faecium* and *E. coli* requires the production of a β-lactam-insensitive D,D-carboxypeptidase to generate the tetrapeptide substrate of LDTs ^11,31^. In contrast, production of the essential LDT-tetrapeptide substrate by a β-lactam-sensitive D,D-carboxypeptidase in *C. difficile* results in the suppression of LDT activity in the presence of the antibiotics. Altogether, our data support the hypothesis that at least one D,D-carboxypeptidase, controlling substrate availability for LDTs, is the primary target for cephalosporins and carbapenems in *C. difficile*. Since 3→3 cross-links are essential for viability in this bacterium ^16^, substrate deprivation for LDTs most likely represents the original mechanism by which cephalosporins and carbapenems impact *C. difficile* survival. Future research will seek to identify the D,D-carboxypeptidase involved, as this enzyme represents an attractive target for the development of new therapies to fight *C. difficile* infections.

## Acknowledgments

We are grateful to Alain Guillot (Micalis, INRAE) and François Fenaille (CEA, Université Paris-Saclay) for helpful discussions. We acknowledge ChemSyBio team (Micalis, INRAE, Jouy-en-Josas) for access to LC-MS/MS facilities. The present work has benefited from Imagerie-Gif core facility supported by l’Agence Nationale de la Recherche (ANR-11-EQPX-0029/Morphoscope, ANR-10-INBS-04/FranceBioImaging; ANR-11-IDEX-0003-02/ Saclay Plant Sciences). This work was funded by the French National Research Agency (ANR-20-CE15-0003-DIFFICROSS to Dr. Johann Peltier).

## Author contributions

Conceptualization, A.O.P., M-P.C-C. and J.P; Methodology, A.O.P. and J.P.; Investigation, A.O.P., G.C. and P.C.; Writing-Original Draft, A.O.P. and J.P.; Writing–Review & Editing, A.O.P., O.S., M-P.C-C. and J.P.; Visualization, A.O.P., P.C., M-P.C-C. and J.P. Supervision A.O.P. and J.P.; Funding Acquisition, J.P.

## Supplemental information titles and legends

Table S1. Excel file containing additional data too large to fit in a pdf, related to Figures 1, 4, S3, S6 and S7.

Document S1. Figures S1-S7 and Tables S2-S6.

## STAR Methods

### Resource availability

#### Lead contact

Further information and requests for resources and reagents should be directed to and will be fulfilled by the lead contact, Johann Peltier.

#### Materials availability

Strains generated in this study are available from the lead contact without restrictions.

#### Data and code availability

- LC-MS/MS datasets have been deposited in the GLYCOPOST repository (GPST000426) and are publicly available as of the date of publication.. RNA-seq and whole genome sequencing data have been deposited to the NCBI Sequence Read Archive (SRA) BioProject PRJNA1107170 and are publicly available as of the date of publication.
- This paper does not report original code.
- Any additional information required to reanalyze the data reported in this paper is available from the lead contact upon request.

### Experimental models and study participant details

#### Bacterial strains and growth conditions

*C. difficile* strains were grown in an anaerobic Jacomex workstation with an atmosphere of 5% H_2_, 5% CO_2_ and 90% N_2_. *C. difficile* strains were cultured in Brain Heart Infusion broth (BHI, BD Difco) or Sporulation Medium for *Clostridium difficile* (SMC, 9% Bacto peptone, 0.5% proteose peptone, 0.15% Tris base, 0.1% ammonium sulphate). All the *C. difficile* strains are described in Table S5.

Plasmids were maintained in *E. coli* strain NEB10β and transformed using standard procedures^35^. *E. coli* HB101 carrying the plasmid RP4 was used for plasmid conjugation with *C. difficile* strains ^36,37^. The *E. coli* strains were cultured in Luria Bertani broth (LB Lennox, Sigma) supplemented with chloramphenicol at 15 µg/mL or 100 µg/mL ampicillin when required, and were grown aerobically at 37°C.

The growth was followed by optical density reading at 600 nm. Growth curves were performed in 96-well plates, with a starting culture at a normalized OD_600_ ≈ 0.05, under anaerobic conditions at 37 °C, in Stratus Plate Reader (Cerillo). The OD_600_ value for each well was recorded every 30 min.

## Method details

### Construction of the C. difficile ldt mutants

All primers used in this study are listed in Table S6. The *C. difficile ldt* deletion mutants were created using a toxin-mediated allele exchange method ^38^. Briefly, ∼800 bp homology arms flanking the region to be deleted were amplified by PCR from *C. difficile* 630Δ*erm*. Purified PCR products were cloned into the PmeI site of the pseudo-suicide allele-coupled exchange (ACE) vector pMSR0 ^38^ using NEBuilder HiFi DNA Assembly (New England Biolabs). The resulting plasmids, listed in Table S5, were transformed into *E. coli* strain NEB10β (New England Biolabs) and all inserts were verified by sequencing. Plasmids were then transformed into *E. coli* HB101(RP4) and transferred by conjugation into the appropriate *C. difficile* strains. Transconjugants were selected on BHI supplemented with *Clostridioides difficile* Selective Supplement (CDSS; Oxoid), and 7.5 μg/mL thiamphenicol. Allelic exchange was performed as described ^38^. All strains were confirmed by locus amplification. The triple mutant was further confirmed by whole-genome sequencing.

### Whole genome sequencing

*C. difficile* strains were grown in BHI for 24h. Cells were harvested and genomic DNA was isolated using NucleoSpin Microbial DNA Mini kit for DNA (Macherey-Nagel). Bacterial genomes were sequenced at Plasmidsaurus (https://www.plasmidsaurus.com/) using the long-read sequencing technology from Oxford Nanopore Technologies (ONT). Using Galaxy Europe, quality controls were performed with FastQC and nanoplot and adapter trimming was performed with Porechop. The resulting reads were mapped to the reference genome using Map with minimap2. Small indels (<50 bp) and SNPs were identified using Clair3, bcftools norm and SnpSift Filter. Large indels were identified using CuteSV. Raw sequence files were deposited to the NCBI Sequence Read Archive (SRA) BioProject PRJNA1107170.

### Fluorescence microscopy

The sample preparation for fluorescence microscopy was carried out under anaerobic conditions. *C. difficile* strains were cultured in BHI. When required, 1 mL culture was incubated with HADA at a final concentration of 10 µM (λ_em_ ≈ 450 nm, Tocris Bioscience) ^19^ for 10 min and washed 3X with anaerobic PBS. Cells were spotted on GeneFrame slides with 1.5% agarose patches. Microscopy was carried out with spinning disk confocal microscopy, using an Inverted Eclipse Ti-E (Nikon) equipped with CSU-X1-A1, Nipkow Spinning Disk confocal system (Yokogawa) and ORCA-Flash4.0 LT CMOS camera (Hamamatsu). Data and statistical analysis were done with MicrobeJ version 5.13I plugin for ImajeJ ^39,40^. Recognition of cells was limited to 1 µm^2^ minimum and 1 - 16 µm length. Cells with defective detection were excluded from analysis. Fluorescent intensity profiles of contour and medial were used for analysis of 2 independent biological replicates. Representative pictures were prepared for publication in CorelDRAW X8 (Corel).

### Spore purification

Overnight cultures of *C. difficile* ΔΔΔ*ldt* and parental strains grown in BHI were spread out on twelve SMC agar plates for sporulation ^41^. After 7 days of anaerobic incubation at 37°C, cells and spores were harvested in 2 ml of ice-cold sterile water. Crude suspensions were washed 10 times with ice-cold sterile water and spores were purified with a HistoDenz (Merck) gradient 20-50%.

### Purification and structural analysis of PG

Vegetative PG was extracted from *C. difficile* strains as previously described with some modifications ^14^. *C. difficile* cultures were grown to OD_600_ ≈ 1.0 at 37°C in BHI with the addition of cefoxitin (64 µg/mL), ceftriaxone (32 µg/mL), meropenem (1 µg/mL) or imipenem (2 µg/mL), when required. Cells were harvested by centrifugation at 5000g for 10 min and processed in a FastPrep apparatus (MP Bioscience) for 30 s at 4 m/s to break the cells. For spore PG extraction, purified spores were additionally incubated twice in decoating buffer (50 mM Tris-HCl pH8.0, 8M Urea, 1% sodium dodecyl sulfate (SDS), 50 mM DTT) for 1h at 37°C. Samples were harvested by centrifugation at 17000g for 10 min.

Both cell and spore pellets were resuspended in cold H_2_O, boiled for 10 min, cooled again, and centrifuged at 17000g for 10 min. The cells were then boiled in 10% SDS, followed by boiling in 4% SDS and washed 10 times with H_2_O. The insoluble material was then treated with pronase for 90 min at 60°C, followed by incubation with DNase, RNase, Lipase and trypsin for 20h at 37°C to purify the cell wall. Finally, the samples were incubated in 48% hydrofluoric acid at 4 °C for 16 h to remove wall polysaccharides. Purified PG was digested with mutanolysin (Sigma), and the soluble muropeptides were reduced with sodium borohydride. Muropeptides were analyzed by LC-MS/MS with an UHPLC instrument (Vanquish Flex, Thermo Scientific) connected to a Q-Exactive Focus mass spectrometer (Thermo Fisher Scientific) fitted with an H-ESI electrospray source (facilities located at ChemSyBio, Micalis, INRAE, Jouy-en-Josas). They were separated by reverse phase chromatography with a ZORBAX Eclipse Plus C18 RRHD column (100 by 2.1mm; particle size, and 1.8μm; Agilent Technologies) at 50°C using 10 mM ammonium formate buffer (pH 4.6) and a 20 min linear gradient from 0 to 20% methanol at a flow rate of 0.3 ml/min. Mass analysis was performed in positive mode with an acquisition range of 380 -1400 *m/z* at resolution 17,500. The Q-Exactive mass spectrometer was operated with capillary voltage at 3.5 kV and a capillary temperature set at 320°C. MS2 was performed in an acquisition range of 160-1600 m/z with an AGC target at 2.10^5^ with HCD collision anode at energy 25. Data were acquired with the Qual Browser suite (Thermo Xcalibur).

Muropeptides were identified from their *m/z* values and MS/MS spectra when required. They were quantified using Skyline open-source software ^42^ following the procedure for small molecule quantification and considering the different adducts and charge states detected for each muropeptide.

The cross-linking index (CI) was calculated according to Glauner^34^ as follows: CI = (1/2 Σdimers + 2/3 Σtrimers)/Σall muropeptides. The relative amount of muropeptides with a certain side chain (X) with a free terminal carboxyl group (acceptor chain) was calculated according to Glauner ^34^ as follows: percentage (X) = (Σmonomers(X) + 1/2Σdimers + 1/3Σtrimers)/Σall muropeptides.

### Minimum inhibitory concentration (MIC) determination

For the determination of the MIC, strains were grown until OD_600_ of 1.0 and inoculated to a starting optical density 0.05 in 200 µL BHI medium supplemented with the required antibiotic range (amoxicillin, oxacillin, cefoxitine, ceftriaxone, imipenem or meropenem) on 96-well plates. Cultures were incubated at 37°C for 24h. MICs for each strain were determined as the lowest concentration without visible growth, at least in three independent experiments.

### RNA sequencing

For total RNA extraction, *C. difficile* strains were grown in 20 mL BHI medium with 32 µg/mL ceftriaxone, until an optical density of 1.0. Total RNA was isolated as previously described ^43^. Quantification of the RNA samples was performed using the Qubit™ RNA High Sensitivity (ThermoScientifc). Samples were sent for RNA sequencing by Novogene Prokaryotic RNA Sequencing services (Novogene). Analysis of the raw FastQ files was performed with Galaxy Pasteur ^44^. Quality control of the raw data was assessed using FastQC and raw data were then subjected to trimming and filtering with AlienTrimmer^45^. Sequencing reads were aligned to *C. difficile* strain 630 genome (NC_009089.1) with the Bowtie2 software ^46^ using default parameters. SARTools DESeq2 was used to perform normalization and differential analysis using values of the 630Δ*erm* wild-type strain as a reference for reporting the expression data of the ΔΔΔ*ldt* strain. Genes were considered differentially expressed if they had a ≥twofold increase or decrease in expression and an adjusted *P* value (*q* value) ≤ 0.05. Assignment of the functional orthologues (K number) and KEGG pathway was performed automatically using BlastKOALA ^47^ and was then manually edited. RNA-seq raw sequence reads were deposited to the NCBI Sequence Read Archive (SRA) BioProject PRJNA1107170.

---

**Figure S1.**
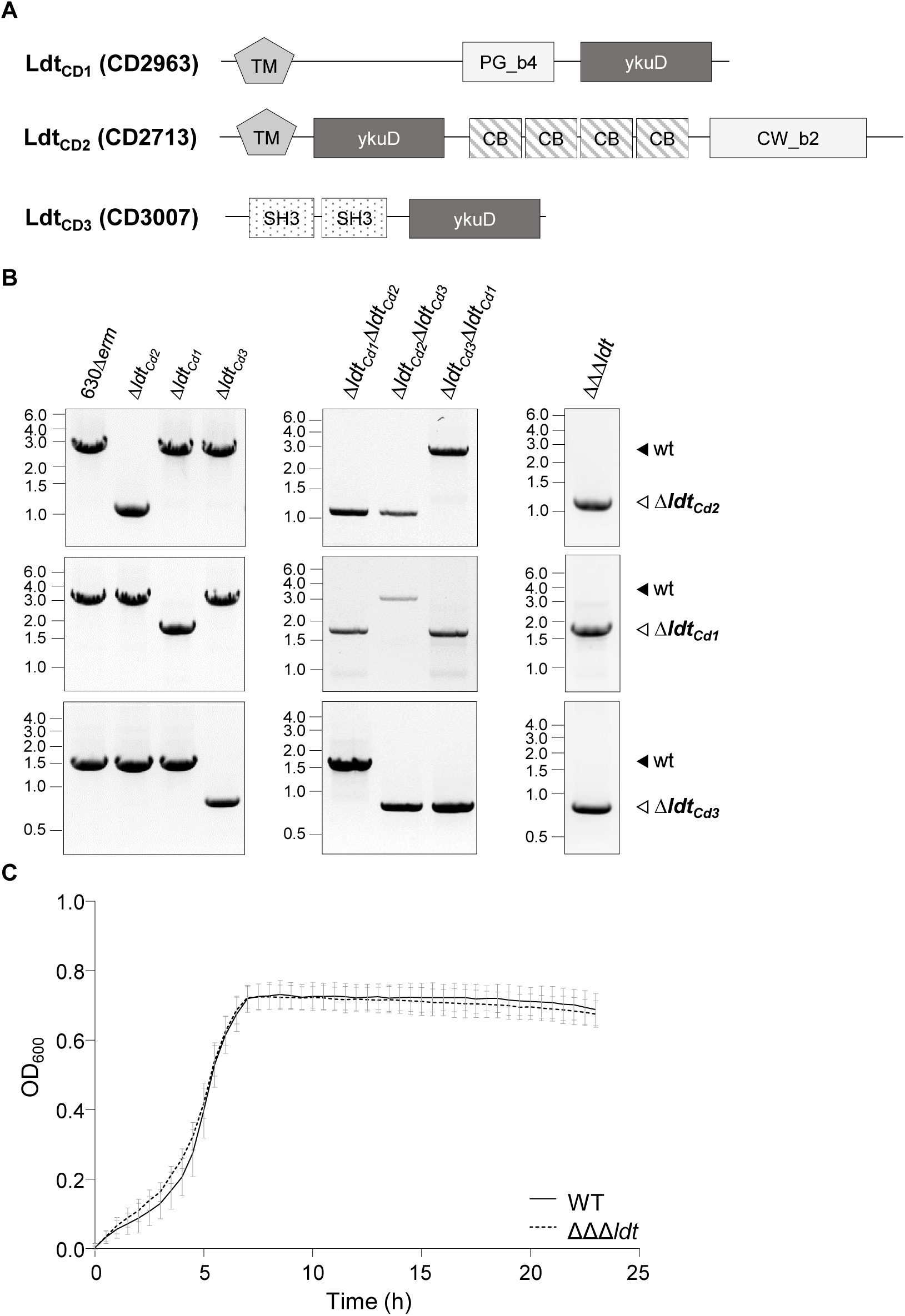
- Impact of a triple mutant of the LDT-encoding genes on *C. difficile* growth. (A) Domain composition of the L.D-transpeptidases Ldt_Cd1_ (CD2963), Ldt_Cd2_ (CD2713) and Ldt_Cd3_ (CD3007) from *C. difficile*. The different conserved domains are represented: transmembrane region (TM in grey), peptidoglycan and cell-wall binding domain (PG_b4 and CW_b2. in light grey), LDT catalytic domain (YkuD in dark grey), cholin-binding domain (CB in stripes) and the SH3 domain associated with a variety of intracellular or membrane-associated proteins (dots). (B) PCR amplification from *C. difficile* 630Δ*erm* wild-type (WT), ΔΔΔ*ldt*, Δ*ldt_Cd1_*, Δ*ldt_Cd2_*, Δ*ldt_Cd3_*, Δ*ldt_Cd1_*Δ*ldt_Cd2_*, Δ*ldt_Cd1_*Δ*ldt_Cd3_* and Δ*ldt_Cd2_*Δ*ldt_Cd3_*, with specific primers to the locus region of *ldt_Cd1_*, *ldt_Cd2_* and *ldt_Cd3_* to verify the different alleles. **C.** Growth curves of *C. difficile* WT and ΔΔΔ*ldt* strains in BHI medium. Means and SD are shown; *n* = 3 independent experiments.

**Figure S2.**
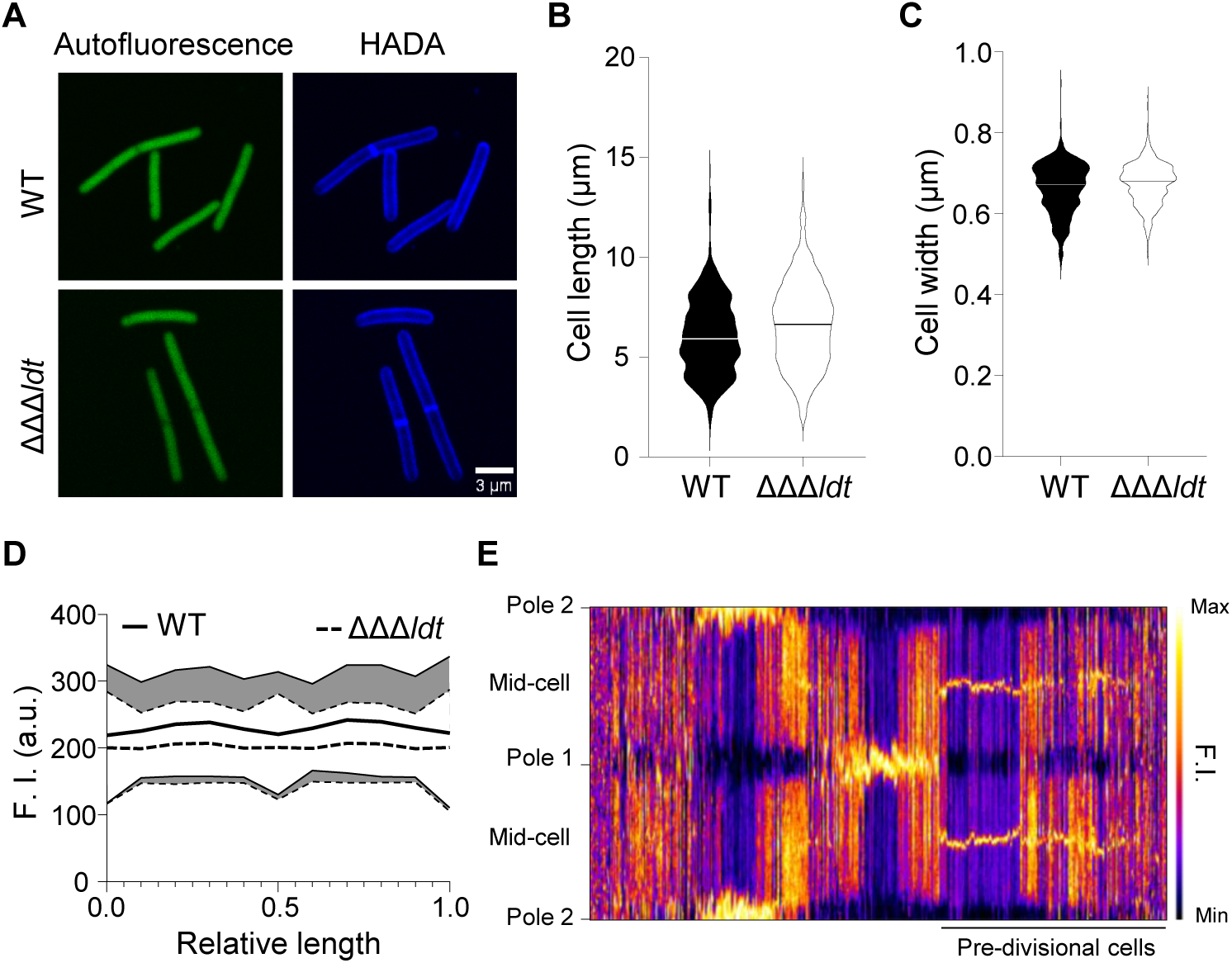
Analysis of cell morphology and HADA incorporation in *C. difficile* 630*Δerm* and ΔΔΔ*ldt*. (A) Cells were collected at an OD_600nm_ of 0.6 and stained with the fluorescent D-amino-acid HADA for 10 min. Cells were imaged on the green channel for the autofluorescence (excitation, 488nm) and on the blue channel for HADA (excitation, 405nm). Scalebar=3 µm. Pictures were treated equally and are representative of 3 independent experiments. (B) Scatter plot showing cell length of 630*Δerm* and ΔΔΔ*ldt* strains with the median of each distribution indicated by a black line. (C) Scatter plot showing cell width of 630*Δerm* and ΔΔΔ*ldt* strains with the median of each distribution indicated by a black line. (D) Average contour fluorescence intensity (HADA distribution). Standard deviation is represented by dashed lines. (E) Analysis of contour fluorescence intensity (HADA distribution) in *C. difficile* 630*Δerm* cells. Cells were grouped by fluorescent distribution and pre-divisional cells were identified. Pole 1 and 2, as well as mid-cell are shown. Scale of fluorescent intensity is depicted. For panels (B) to (E), 2 independent experiments were analysed for *C. difficile* 630*Δerm* (grey, *n*=544) and ΔΔΔ*ldt* (orange, *n*=878).

**Figure S3.**
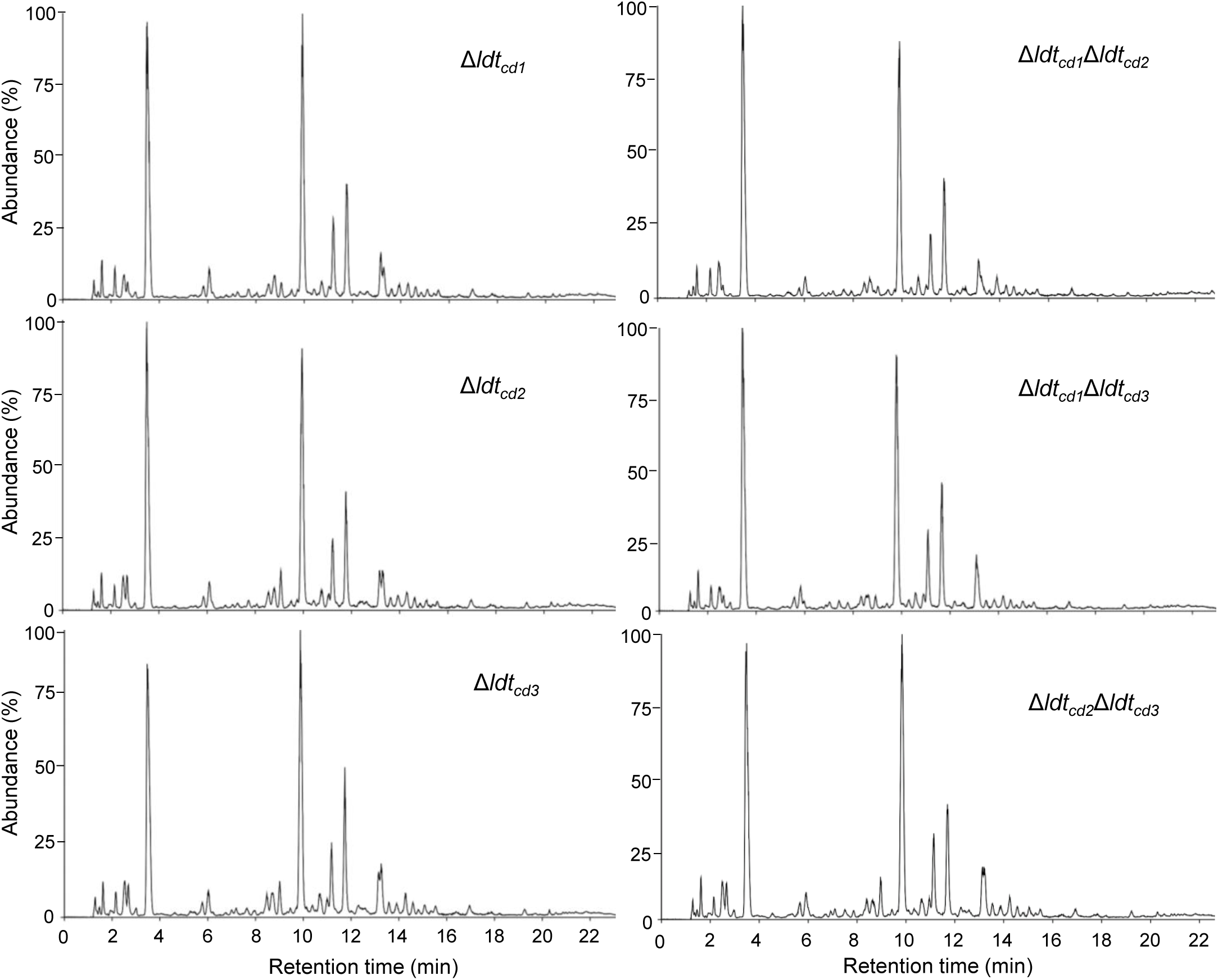
- Impact of *ldt* single and double mutants on the PG structure of *C. difficile* vegetative cells. LC-MS separation of muropeptides from vegetative cells of *C. difficile* Δ*ldt_cd1_*, Δ*ldt_cd2_*, Δ*ldt_cd3_*, Δ*ldt_cd1_*Δ*ldt_cd2_*, Δ*ldt_cd1_*Δ*ldt_cd3_* _and_ Δ*ldt_cd2_*Δ*ldt_cd3_* strains. See also Table S1 for the structure of all identified muropeptides. Data are representative of two independent experiments.

**Figure S4:**
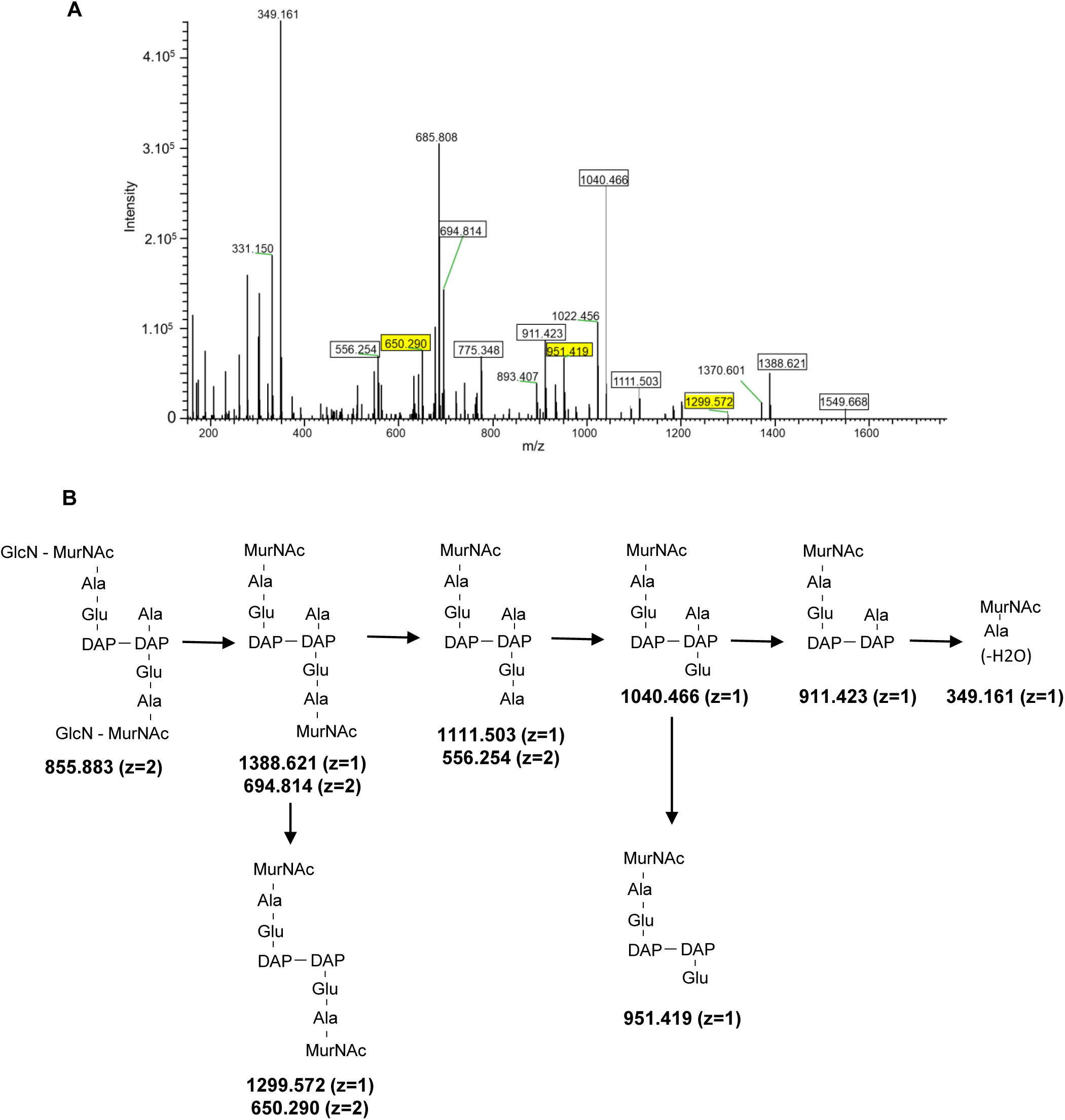
MS/MS analysis of the second muropeptide dimer from the spore cortex of *C. difficile* wild type. (A) MS/MS analysis of the ion (*m/z* = 855.883 with *z*=2) corresponding to the peak with a retention time of 8.75 min in Fig. 4A and Table S2. Fragments (squared) were detected as [M+H]^+^ (z=1) or [M+2H]^2+^ (z=2) adducts. Fragments resulting from loss of a water molecule are not squared. In yellow, the fragments allowing the assignment of the 3-3 cross-link. (B) Structures inferred from the MS/MS analysis are presented. The loss of one alanine from the C-terminal end of different ions (mass loss of 89.048) establishes the presence of a 3→3 crosslink.

**Figure S5.**
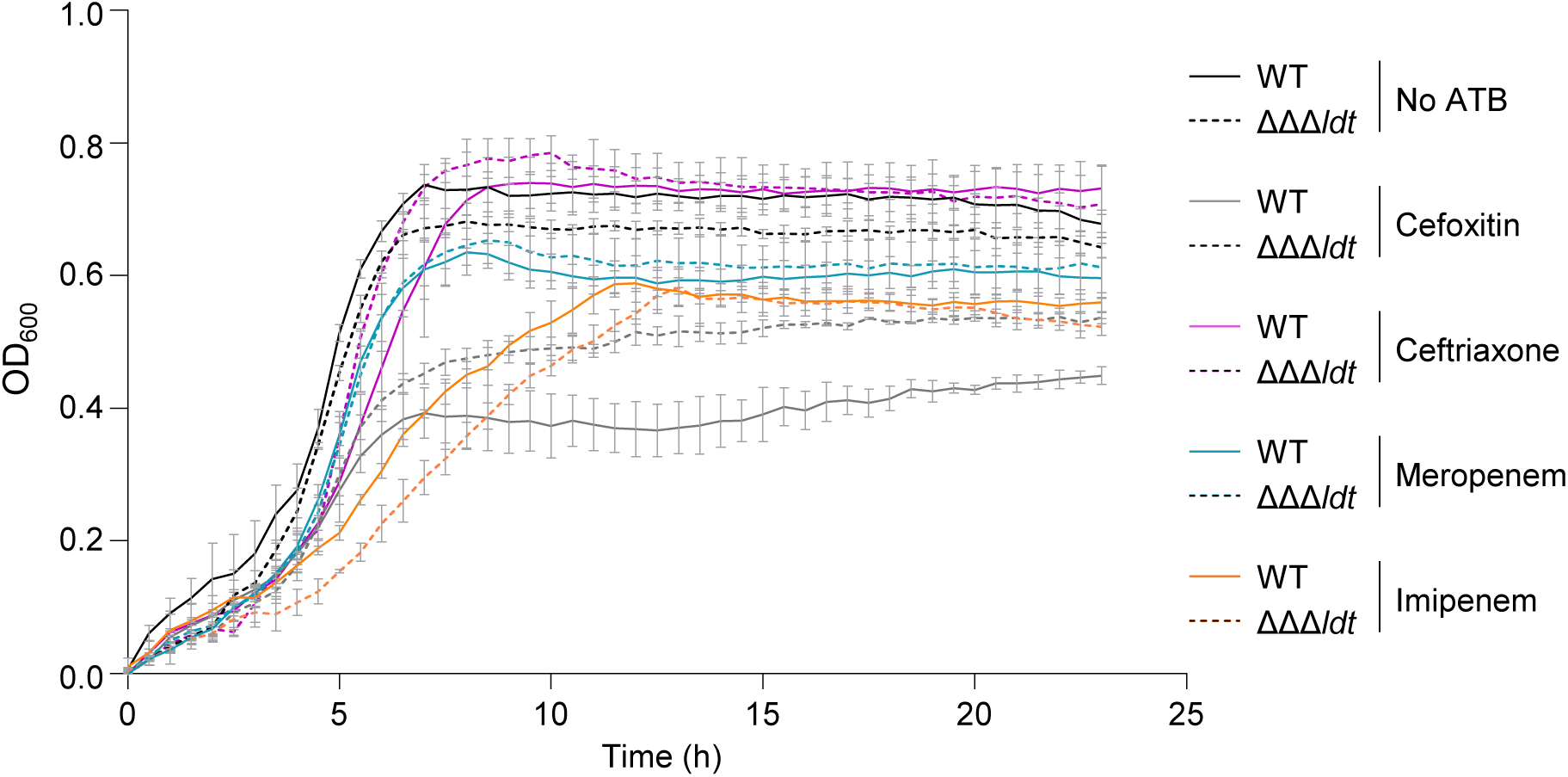
- Growth curve of of *C. difficile* 630Δ*erm* wild-type (WT) and ΔΔΔ*ldt* strains in presence of sub-MIC of β-lactam antibiotics. Means and SD are shown; *n* = 3 independent experiments.

**Figure S6.**
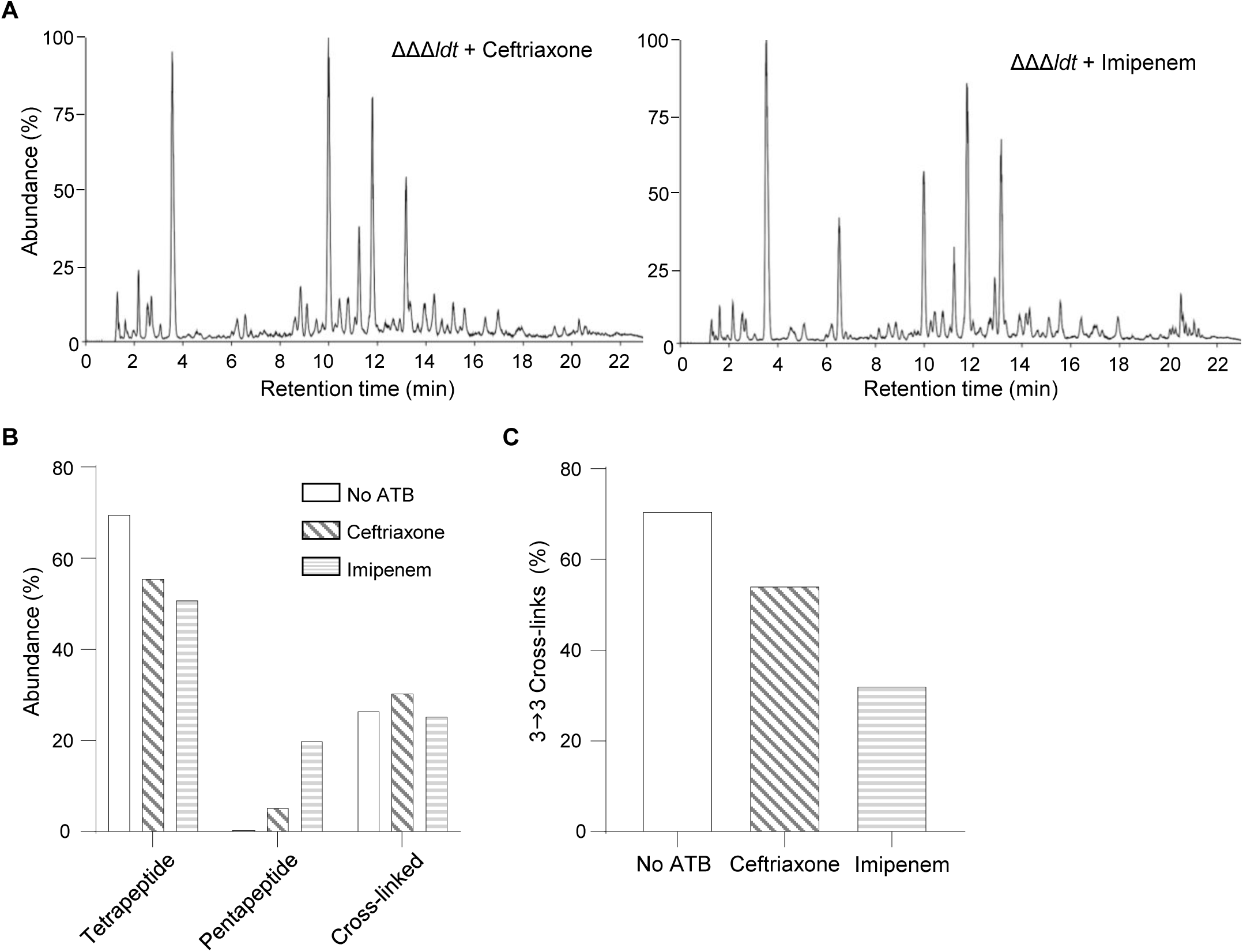
- Impact of the cephalosporin ceftriaxone and the carbapenem imipenem on the PG structure of vegetative cells of *C. difficile* ΔΔΔ*ldt*. (A) LC-MS chromatograms of muropeptides from vegetative cells of *C. difficile* ΔΔΔ*ldt* strain grown in the presence of subinhibitory concentrations of ceftriaxone or imipenem. See also Table S1 for the structure of all identified muropeptides. Data are from one experiment. (B) Abundance of muropeptides with a tetrapeptide or a pentapeptide stem and cross-linking index in the PG of *C. difficile* ΔΔΔ*ldt* strain grown in absence of antibiotics (no ATB) or in presence of ceftriaxone (ceftri.) or imipenem (imip). *n* = 1 experiment. (C) Abundance of muropeptide dimers with a 3→3 crosslink relative to the total crosslinks in dimers in the PG of *C. difficile* ΔΔΔ*ldt* strain grown in the absence of antibiotics (no ATB) or in the presence of ceftriaxone (ceftri.) or imipenem (imip). *n* = 1 experiment.

**Figure S7:**
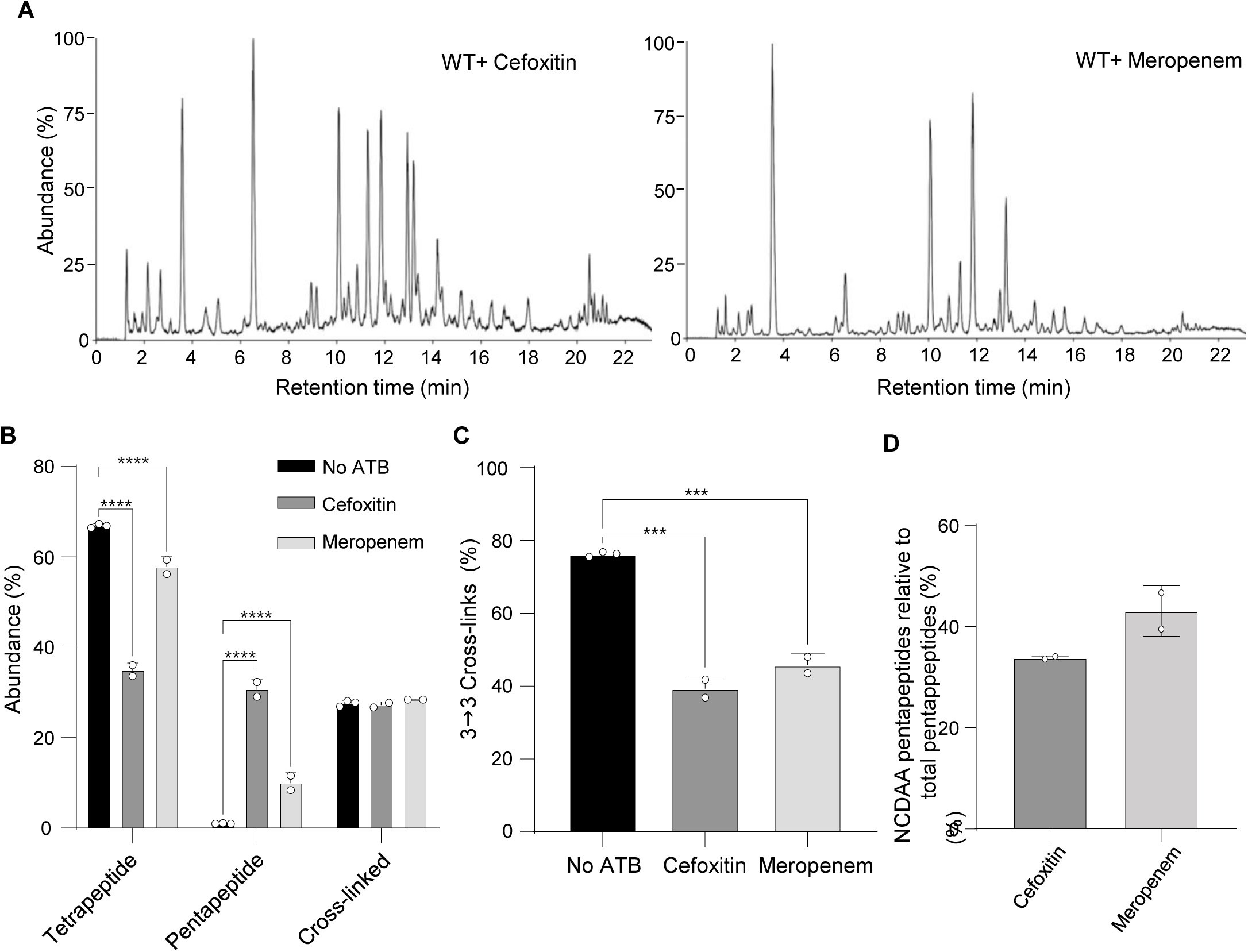
Impact of the cephalosporin cefoxitin and the carbapenem meropenem on the PG structure of vegetative cells of *C. difficile* 630Δ*erm* wild-type (WT). (A) LC-MS chromatograms of muropeptides from vegetative cells of *C. difficile* WT strain grown in the presence of subinhibitory concentrations of cefoxitin or meropenem. See also Table S1 for the structure of all identified muropeptides. Data are representative of two independent experiments. (B) Abundance of muropeptides with a tetrapeptide or a pentapeptide stem and cross-linking index in the PG of *C. difficile* WT strain grown in the absence of antibiotics (no ATB) or in the presence of cefoxitin (cefox.) or meropenem (merop). (C) Abundance of muropeptide dimers with a 3→3 crosslink relative to the total crosslinks in dimers in the PG of *C. difficile* WT strain grown in the absence of antibiotics (no ATB) or in the presence of cefoxitin (cefox.) or meropenem (merop). (D) Abundance of muropeptides with a pentapeptide stem ending with a non-canonical D-amino-acid (NCDAA: Gly, Phe, Leu or Val) relative to the total pentapeptide muropeptides in the WT strain. All graphs represent mean ± SD and include individual data points; *n* = 2 independent experiments. \*\*\**P* ≤ 0.001 and \*\*\*\**P* ≤ 0.001 by a two-way ANOVA followed by a Dunnett’s multiple comparisons test (B and C).

